# Utilizing a Cell-free Protein Synthesis Platform for Natural Product Synthesis

**DOI:** 10.1101/2022.07.22.501086

**Authors:** Alex Ditzel, Fanglong Zhao, Xue Gao, George N. Phillips

## Abstract

Natural products are a great source of pharmaceuticals, providing a majority of all small molecule drugs that exist today. However, creating natural products through organic synthesis or in heterologous hosts can be difficult and time-consuming. Therefore, to allow for easier screening and production of natural products, we demonstrated the use of a cell-free protein synthesis (CFPS) system to partially assemble natural products *in vitro* using coupled enzyme reactions. The tea caffeine synthase TCS1 was utilized to synthesize caffeine within a CFPS system. Cell-free systems also provide the benefit of allowing the use of substrates that would normally be toxic in a cellular environment to synthesize novel products. The automation and reduced metabolic engineering requirements of CFPS systems combined with other synthesis methods can allow for the efficient generation of new compounds.

## Introduction

Natural products are an essential and highly varied category of small molecules, defined as being produced naturally by some organism, and these compounds are extremely important to the pharmaceutical industry ^1,2^. In fact, the majority of all small-molecule testing in the pharmaceutical industry makes use of natural products and they are considered to be the most productive source of new drugs ^2,3^. Despite that fact, many of these companies decommissioned their small molecule discovery programs in favor of combinatorial chemistry due to its ability to create large libraries which can be efficiently screened for useful biological activity ^4^. Unfortunately, these new programs failed to create many new useful compounds, with only a few out of millions having any degree of desirable activity ^5^.

The reason pharmaceutical companies seek out natural products is because of their inherent biological activities. Natural products can be, however, quite large and complex. For example, two large, valuable groups of natural products are based on polyketides, such as erythromycin, and nonribosomal peptides, such as daptomycin ^6^. Additionally, Taxol, a diterpene that is perhaps the most important anti-cancer therapeutic in history, is a natural product derived from yew tree bark ^7^. Taxol is used for the treatment of lung, breast, prostate, bladder, ovarian, esophageal, and other many other types of cancer and is credited with saving countless lives since it was approved for use in 1993 ^8^. Current methods of producing Taxol include biosynthesis of precursors in cell culture followed by organic synthesis to complete the process ^9^. It seems likely that a combination of biosynthetic and organic synthesis will be most practical for the generation of many natural products and natural product-like compounds.

For a biosynthetic approach, the relevant enzymes and pathways need to be identified. The conventional method for producing enzymes from bacteria involves transforming cells so that they contain a plasmid with a gene encoding the desired product and growing large numbers of those cells ^10^. However, in recent years it has become possible to synthesize proteins *in vitro* by taking advantage of cellular machinery in an open system^11^. Cell-free protein synthesis (CFPS) involves a single tube reaction which goes from plasmid to protein ^11^.

CFPS works by utilizing cellular lysates which contain all the machinery necessary for transcription and translation, such as ribosomes, T7 RNA polymerase, etc. along with additional NTPs as energy sources, so that if a plasmid is introduced mRNA and then protein will be produced. T7 polymerase is used to provide specificity to plasmids containing the corresponding promoter sequence ^12^. CFPS systems most commonly use an *E. coli* based system, but wheat germ-based systems have also been used to synthesize a wider range of eukaryotic proteins^13^. Additionally, yields can be improved through the introduction of a feeding buffer containing NTPs and other substrates, as well as through continuous flow-based systems that continually replenish substrate ^14^.

There are several advantages to the use of CFPS as opposed to *in vivo* production. One important advantage is the capability to produce and utilize toxic compounds, due to the lack of cells that would be killed by cytotoxic effects, expanding the range of compounds that can be produced by the system. Another advantage is that the process can be fully automated from the introduction of the plasmid into the system to the production of the protein ^15^. Additionally, the need for metabolic engineering is greatly reduced because the metabolic load can be essentially ignored. Finally, the speed of reactions in these systems is also significantly faster, taking only 1 day to go from plasmid/substrate to pure protein/product^16^.

As part of the path to realize these advantages, a CFPS-based system has been used to perform the function of the first two of five modules involved in the synthesis of the nonribosomal peptide gramicidin S, a valuable antibiotic ^17^. Relatively high yields of approximately 12 mg/mL for the production of the natural product gramicidin S were obtained using this method, higher than existing cell-based systems ^17^. Cell-free systems have also been used to incorporate novel molecules, such as non-standard amino acids into a system in ways that would be more difficult *in vivo* ^18^. However, the potential to utilize these systems to synthesize natural products from plasmids and precursors in a single reaction has not been fully explored. We extend this research by working to produce different compounds and natural products using a CFPS system as shown in Figure 1.

**Figure 1.**
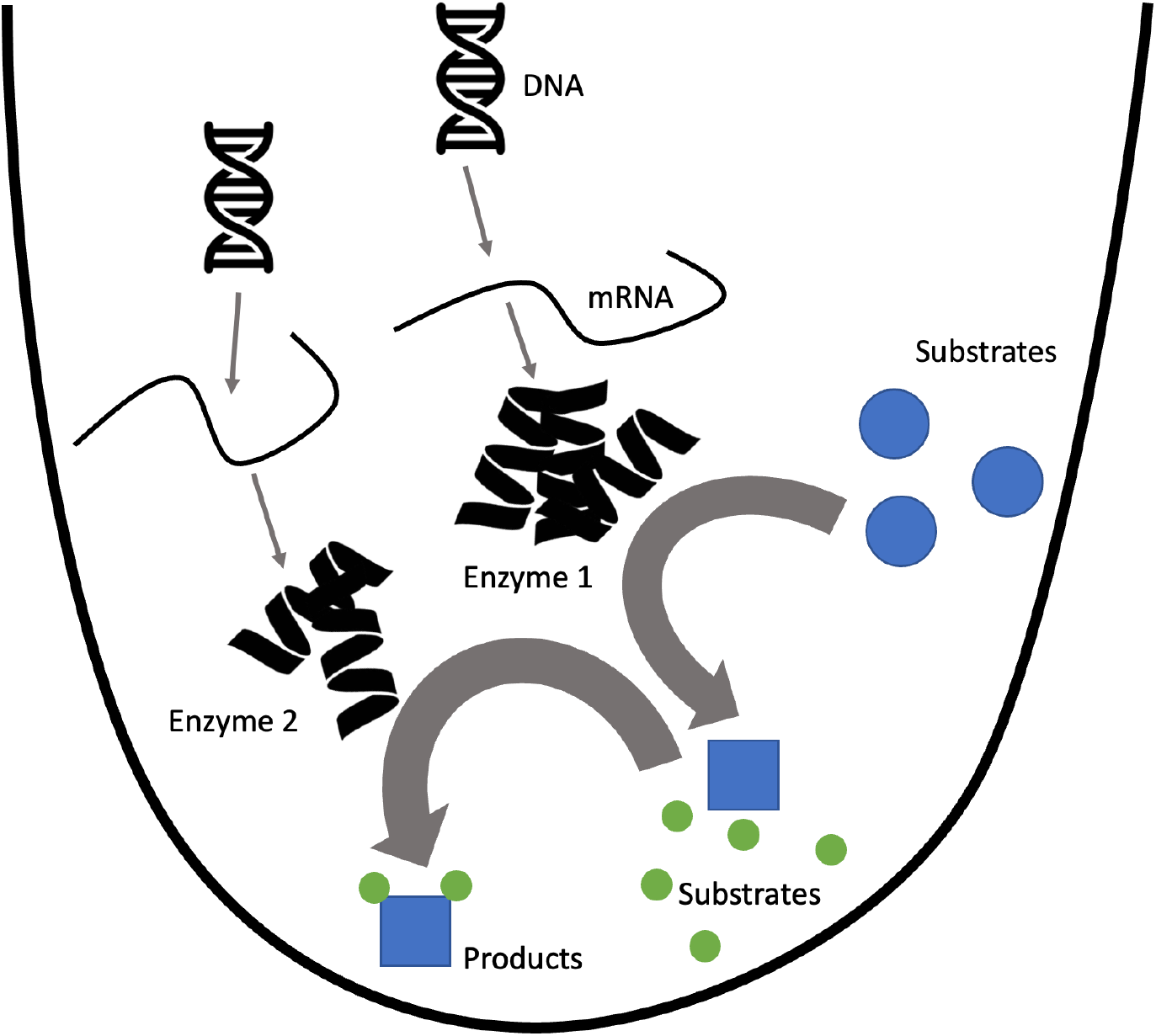
A simplified view of natural product synthesis in a cell-free system. The system is shown with introduced DNA which is then transcribed to mRNA and translated to proteins which then act on substrates within a single solution.

To further show the utility of CFPS systems in natural product synthesis, we chose methylation, a common reaction involved in the synthesis of many natural products, and in particular methylation performed by SAM-dependent methyltransferases (MTases). This is a fairly broad category of enzymes that perform methylation reactions on many different substrates. They also provide the possibility for future work with the application of alkylation activity of some SAM-dependent MTases in the presence of SAM analogues ^19,20^. The promiscuity of these enzymes to utilize SAM analogues to transfer alkyl groups can be difficult to utilize due to the toxicity of many SAM analogues in cells, but in a CFPS their activity could be used more easily.

We chose the synthesis of caffeine as an example of a common and useful natural product that is created by SAM-dependent MTases. There are already established methods for caffeine purification and identification using LC-MS, and the enzyme TCS1 (tea caffeine synthase) has previously been expressed in *E. coli* and found to express well and be soluble^21^. These advantages made caffeine seem like a suitable choice for further application of using CFPS systems for product synthesis.

Caffeine can be described as a tri-methylated form of xanthine and has the alternative name 1,3,7-trimethyl xanthine, which fits with how the compound is synthesized in nature. Typically, caffeine is made in multiple methylation steps going from xanthine or xanthosine to a mono-methylated xanthine, then a di-methylated xanthine, and finally to caffeine^22,23^. In this paper, we are using the tea caffeine synthase enzyme, TCS1, which performs 2 different methylation steps in the production of caffeine in tea plants, going from 7-methylxanthine to theobromine to caffeine^24^. We will be utilizing its activity in methylating theobromine to caffeine in a CFPS system in this study. (Figure 2)

**Figure 2.**
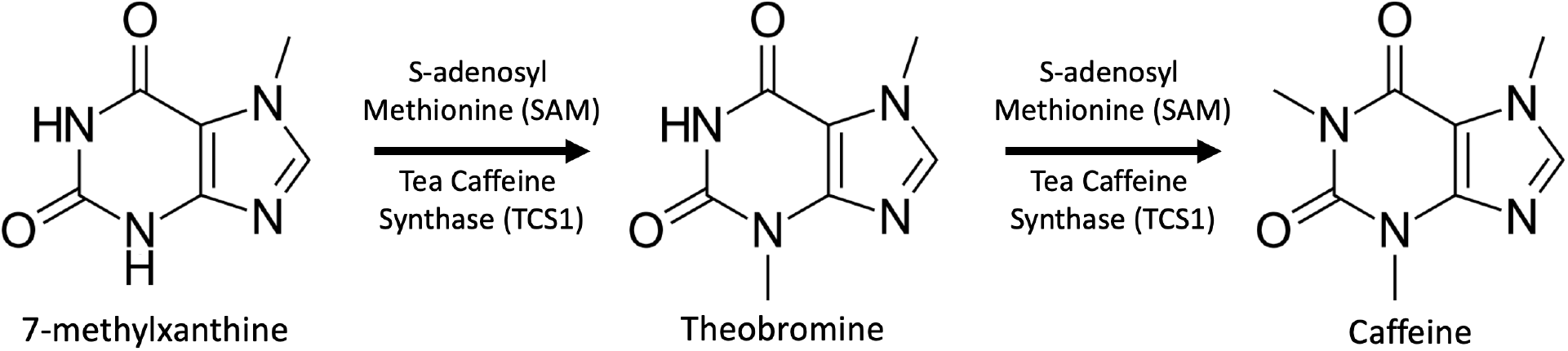
TCS1 Caffeine production pathway in this study. TCS1 produces caffeine by the methylation of 7-methylxanthine to theobromine and then to caffeine.

## Results and Discussion

CFPS systems have immense potential as controlled systems for the synthesis of small molecules. To actualize that potential, we show the production of a natural product within a CFPS system. These synthetic systems, together with the provision of substrates, have the potential to synthesize products more quickly for rapid prototyping and testing natural products. To demonstrate this potential, caffeine was chosen as a product that is useful and can be produced by an enzyme, TCS1, which has been previously shown to be produced in *E. coli*. Therefore, the production of TCS1 within a CFPS system was demonstrated and then the system was further used to synthesize caffeine as described below.

To demonstrate the production of caffeine in the cell-free system, we performed several intermediate experiments. We first synthesized the enzymes of interest using the TCS1 gene in the CFPS system. On an SDS-PAGE gel, the apparent mass of the protein seemed to match the expected mass for the TCS1 enzyme and was relatively pure, without other bands visible (Figure 3). Therefore, the CFPS system seemed to be successful in synthesizing TCS1 protein. To demonstrate activity, we also performed caffeine production assays.

**Figure 3.**
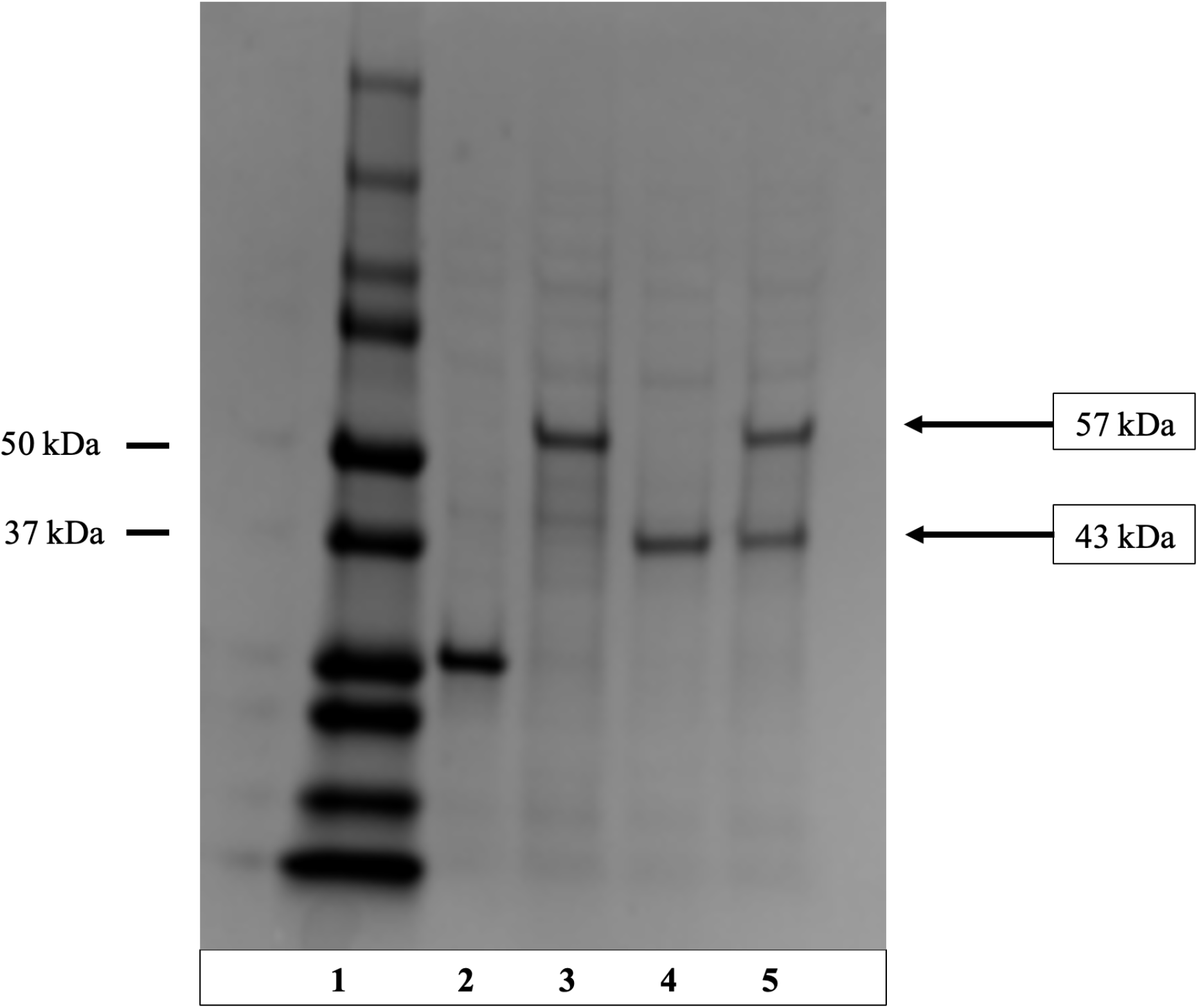
Production of GUD1, a guanine deaminase which can convert guanine to xanthine in the early stages of caffeine biosynthesis, and TCS1 in a CFPS system. An SDS-PAGE gel shows the production of Guanine Deaminase (GUD1) and the tea caffeine synthase TCS1 from the CFPS system. Lane 1 is the ladder. Lane 2 is a GFP control. Lane 3 is GUD1 (guanine deaminase from *S. Cerevisiae*) with an MW of ∼57 kDa. Lane 4 is TCS1 with an MW of ∼43 kDa. Lane 5 is GUD1 and TCS1 produced together. This demonstrated the capability of the system to synthesize multiple enzymes at once without major issues.

We assayed the methylation activity of TCS1 by taking the purified enzyme from the CFPS system and performed a methylation reaction with both theophylline and theobromine substrates. When the TCS1 protein synthesized with the CFPS system was incubated with theophylline and SAM, there were no significant peaks observable by LC-MS other than the substrates (Figure 4B). However, when the TCS1 was incubated with theobromine and SAM there was a peak observable at the same retention time and mass as the pure caffeine standard (Figure 4C). This demonstrates that the TCS1 produced in the CFPS system was functional and it was able to synthesize caffeine. Additionally, despite some previous research suggesting that theophylline may be an adequate substrate for caffeine production using TCS1^21^, we found that it did not result in any caffeine production in the conditions we were using, but theobromine worked well and resulted in easily visible caffeine peaks when checked with LC-MS as seen in figure 4.

**Figure 4.**
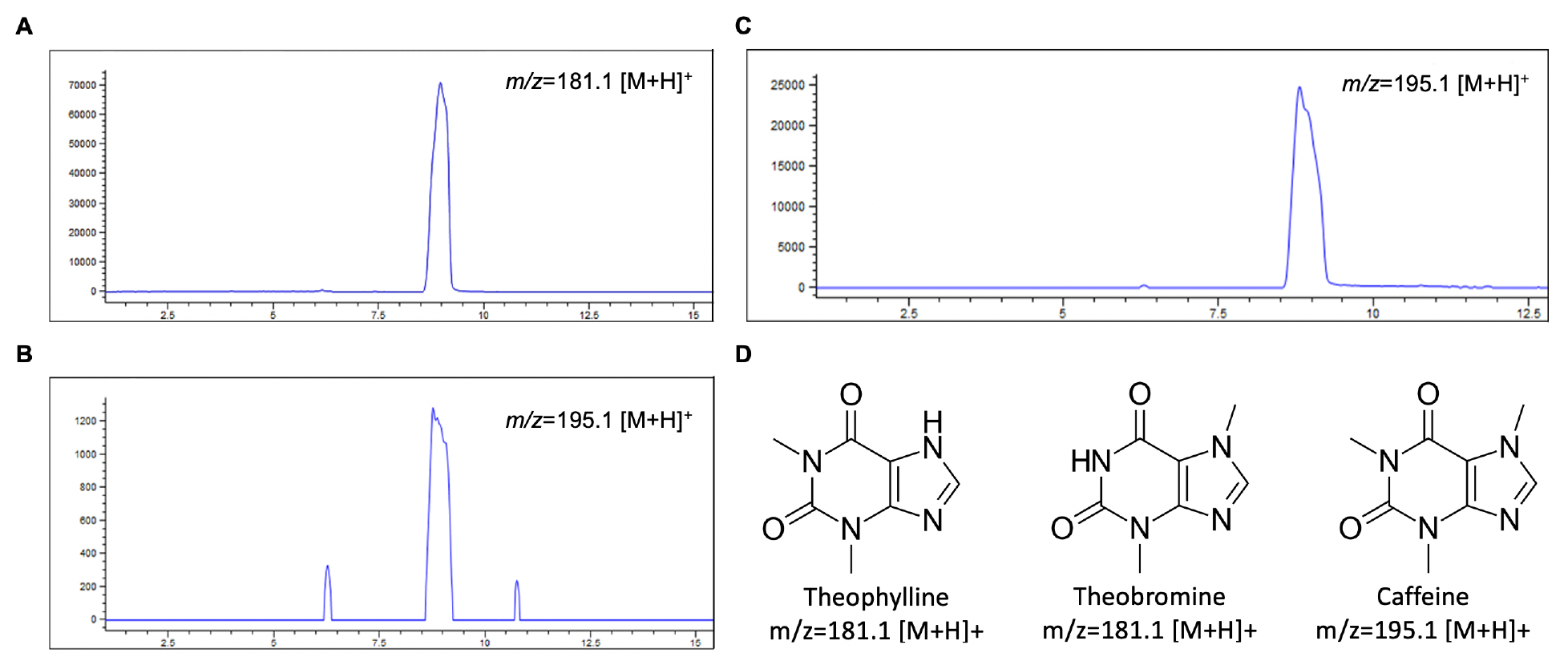
Production of caffeine from theophylline and theobromine substrates using purified TCS1. LC-MS chromatograms show the presence of product or reactant after chloroform extraction from the reaction mixture. The Y-axis is intensity in arbitrary units and the X-axis is retention time on the column in minutes. A) Mass filter analyses of the theophylline substrate (m/z=181.1 [M+H]^+^) B) Mass filter analyses of the enzymatic reaction of TCS1 and substrate theophylline. The final product substrate caffeine (m/z=195.1 [M+H]^+^) can be detected in the reaction, but the intensity here is close to the level of noise, showing very little, if any, product was made. C) With a theobromine substrate mass filter m/z=195.1 [M+H]^+^ is used to detect caffeine, and the product is clearly produced with detectable yield due to the clear peak at the expected mass. D) The structures of theophylline, theobromine, and caffeine are shown along with their expected m/z in LC-MS chromatograms

To test the ability of TCS1 to make caffeine within the CFPS system without purification, the SAM and theobromine substrates were supplied in addition to the other compounds necessary for the CFPS system. When the resulting solution was checked with LC-MS after incubation, there were several peaks present due to the variety of compounds within the CFPS system (Figure 5A). However, there was one additional peak that was only present when theobromine and SAM were added, with the m/z 195.1 [M+H]^+^and the same retention time as the pure caffeine standard (Figure 5). Quantification of the amount of caffeine lacked accuracy due to inefficiencies in the chloroform extraction process. The presence of caffeine in the final solution demonstrates the simultaneous production of TCS1 and the synthesis of caffeine within the bulk CFPS system.

**Figure 5.**
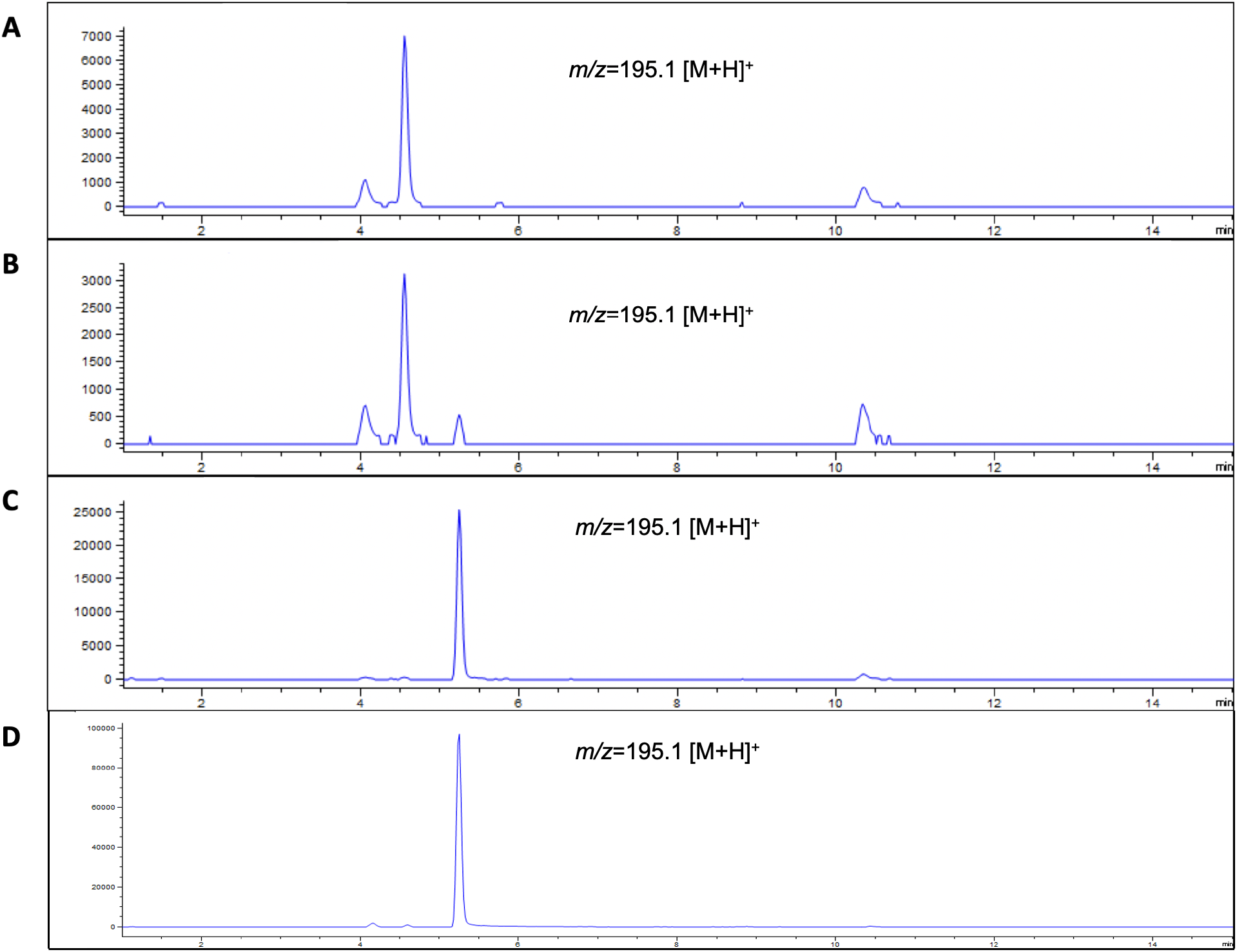
Production of caffeine within a CFPS system. LC-MS chromatograms showing the presence of product or reactant after chloroform extraction from the reaction mixture. The Y-axis is intensity in arbitrary units and the X-axis is retention time on the column in minutes. Each chromatogram is shown with m/z=195.1 [M+H]^+^which would show the presence of caffeine. A) A negative control chromatogram is shown which is the result of a chloroform extraction of the CFPS system producing TCS1 without the presence of theobromine substrate added. B) The experimental chromatogram which shows the result of a chloroform extraction on a CFPS system that had theobromine added and simultaneously produced TCS1. C) A positive control chromatogram which shows the results of concentrated caffeine after a chloroform extraction to show the location of the caffeine peak. D) A caffeine standard chromatogram to show the expected characteristic peak for caffeine

The experiments in the CFPS system with TCS1 demonstrate the possibility of using CFPS systems more generally as a platform for natural product synthesis. Furthermore, this result shows that CFPS systems can be used for single-solution reactions that can both synthesize an enzyme and utilize that enzyme for product synthesis in one simple process without requiring enzyme purification.

The prospect of utilizing CFPS systems as a platform for natural product synthesis is very exciting as it has the potential to provide several benefits over conventional synthesis methods. As demonstrated here with caffeine, it is possible to utilize a single-step mixture of a plasmid containing the gene of interest, the substrate, and any necessary cofactors within the CFPS system without the complication of multiple steps of protein synthesis and running the reactions separately. In this case, the simplified process may be useful for a workflow that involves rapid prototyping by synthesizing a small amount of a variety of different compounds for screening. Additionally, while the methylation of theobromine to make caffeine was the reaction of interest here, other SAM-dependent methylation reactions will likely behave similarly in CFPS systems and thus CFPS systems could be used for a variety of methylation reactions.

One particularly useful advantage of cell-free systems over cellular systems is their resilience against compounds which are toxic to cells, but which have a mechanism of toxicity that does not directly interfere with the limited reactions that occur within a CFPS system. This provides the prospect of utilizing CFPS systems in conjunction with substrates that would normally be toxic to cells to synthesize products that could not normally be synthesized in cells. For example, expanding upon the use of SAM-dependent methylation in this paper, it may be possible to utilize S-adenosyl ethionine or other alkyl analogues of SAM to synthesize a variety of alkylated products that would not normally be able to be made in cells. This creates the potential to create a variety of new products based on existing natural compounds which cannot be synthesized easily within cells. In the future, we hope to demonstrate the utility of CFPS systems for small molecule synthesis and show that the system can be used to create novel products which could not be made within conventional cellular systems.

## Methods

### Plasmid Preparation

The plasmid containing the *TCS1* gene was made by performing a codon optimization on the tea caffeine synthase gene so that it would be suitable to expression efficiently in *E. coli* with the IDT codon optimization tool. This optimized gene was then ordered as a gBlock from IDT and inserted into the *pNIC28* plasmid through the NEBuilder homology-based assembly method. This plasmid was then transformed into the DH5α *E. coli* strain to generate more plasmid, which was then used in the CFPS system to synthesize the TCS1 protein.

### Cell-free Protein Synthesis and Purification

The cell-free protein synthesis was performed using a Bioneer Exiprogen system using *E. coli* cellular lysate and Bioneer’s provided master mix of reagents. The reaction was performed in a Exiprogen Tag-free system which includes TEV protease to cleave the TEV site which was used to attach the 6xHIS tag to the TCS1. The reaction was performed at 30° C using the standard Exiprogen Tag-free protocol as recommended by Bioneer. The protein was then purified with Ni beads and the mass of the protein was checked on an SDS-PAGE gel.

### Caffeine synthesis using purified TCS1 from CFPS system

To verify the function of the TCS1, TCS1 was synthesized using a CFPS system, purified, and the 6x HIS tag was cleaved. This TCS1 protein was then used to perform an *in vitro* reaction with TCS1 in a pH 8.0 0.1 M Tris solution, along with 200 μM MgCl_2_, 50 μM SAM, and 200 μM theobromine. The products were then extracted from this system by performing a chloroform extraction and the products were checked using LC-MS to verify the presence of caffeine.

### Caffeine Synthesis and Extraction from CFPS System

To perform the caffeine synthesis within the CFPS system the substrate theobromine and the cofactor SAM were added to the CFPS system along with the plasmid containing the gene for TCS1. After the protein was synthesized, a chloroform extraction was performed on the CFPS system to remove the caffeine from the system in a form that was more easily analyzed. Chloroform was added to a part of the system and mixed with the aqueous solution. Then the chloroform layer was drawn off using a glass pipette and the chloroform was evaporated to leave the caffeine in a purer form for analysis with LC-MS.

### Liquid Chromatography Mass Spectrometry Protocol

To verify the production of caffeine from the TCS1 LC-MS was used. The analysis was achieved by LC-UV-MS analysis on an Agilent Technologies 6120 Quadrupole LC-MS (with UV-detector). Agilent Eclipse Plus C_18_ column (4.6×100 mm) was used for metabolites separation with a linear gradient of 5-95% acetonitrile (v/v) over 15 min in H_2_O (0.1% formic acid, v/v) at a flow rate of 0.5 mL/min.

## Supporting information

Supplemental figures, DNA sequence, and computational modeling work

## Author Contributions

A.D. designed and performed experiments, wrote and edited manuscript.

F.Z. performed LC-MS experiments and revised the manuscript.

X.G. supervised analysis experiments and revised the manuscript.

G.P. designed experiments, wrote and edited manuscript.

## Acknowledgement

Supported by a training fellowship from the Gulf Coast Consortia, on the Houston Area Molecular Biophysics Program (Grant No. T32 GM008280) to A.D., by NIH grants R01 GM115261 and R01 CA217255 to G.P., and by the NIH grant R35 GM138207 and the Robert A. Welch Foundation (C-1952) to X.G.

