## Supplemental figures, DNA sequence, and computational modeling work for "Utilizing a Cell-free Protein Synthesis Platform for Natural Product Synthesis"

DNA sequence of TCS1 gene used:

ATGGAACTGGCTACCGCTGGTAAAGTTAACGAAGTTCTGTTCATGAACCGTGGTGAAGGTGAATCTTCT
TACGCTCAGAACTCTTCTTTCACCCAGCAGGTTGCTTCTATGGCTCAGCCGGCTCTGGAAAACGCTGTTG
AAACCCTGTTCTCTCGTGACTTCCACCTGCAGGCTCTGAACGCTGCTGACCTGGGTTGCGCTGCTGGTCC
GAACACCTTCGCTGTTATCTCTACCATCAAACGTATGATGGAAAAAAAATGCCGTGAACTGAACTGCCAG
ACCCTCGAACTCCAGGTCTACCTGAACGACCTGTTCGGTAACGACTTCAACACCCTGTTCAAAGGTCTGT
CTTCTGAAGTTATCGGTAACAAATGCGAAGAAGTTCCGTGCTACGTTATGGGTGTTCCGGGTTCTTTCCA
CGGTCGTCTGTTCCCGCGTAACTCTCTGCACCTGGTTCACTCTTCTTACTCTGTTCACTGGCTGACCCAGG
CTCCGAAAGGTCTGACCTCTCGTGAAGGTCTGGCTCTGAACAAAGGTAAAATCTACATCTCTAAAACCTC
TCCGCCGGTTGTTCGTGAAGCTTACCTGTCTCAGTTCCACGAAGACTTCACCATGTTCCTGAACGCTCGTT
CTCAGGAAGTTGTTCCGAACGGTTGCATGGTTCTGATCCTGCGTGGTCGTCAGTGCTCTGACCCGTCTGA
CATGCAGTCTTGCTTCACCTGGGAACTGCTGGCTATGGCTATCGCTGAACTGGTTTCTCAGGGTCTGATC
GACGAAGACAAACTGGACACCTTCAACATCCCGTCTTACTTCGCTTCTCTGGAAGAAGTTAAAGACATCG
TTGAACGTGACGGCAGCTTCACCATCGACCATATAGAGGGCTTCGACCTGGACTCGGTTGAAATGCAGG
AAAACGACAAATGGGTTCGTGGTGAAAAATTCACCAAAGTTGTTCGTGCTTTCACCGAACCGATCATCTC
TAACCAGTTCGGTCCGGAAATCATGGACAAACTGTACGACAAATTCACCCACATCGTTGTTTCTGACCTG
GAAGCTAAACTGCCGAAAACCACCTCTATCATCCTGGTTCTGTCTAAAATCGACGGTTAG

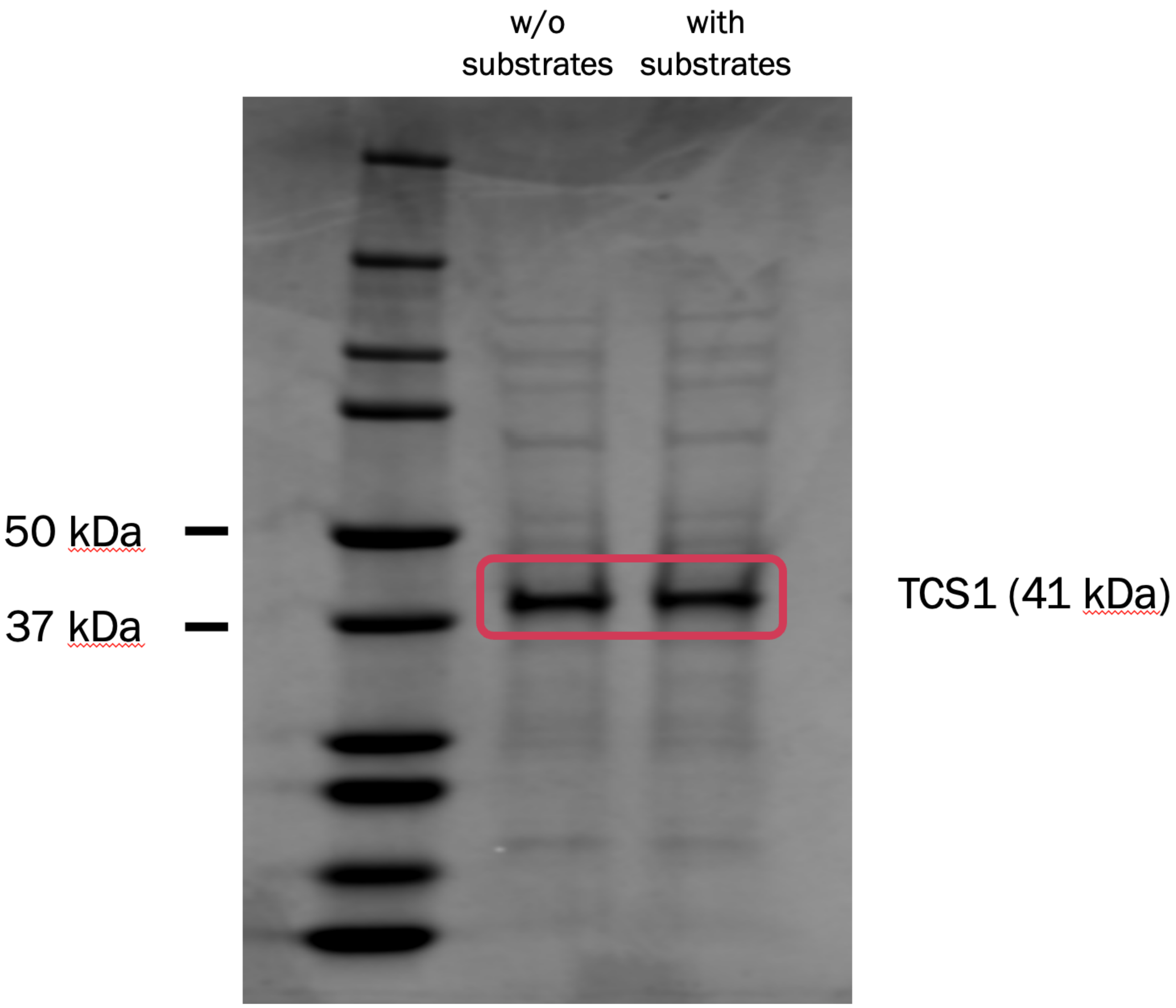


**Figure S1**. Production of TCS1 in a CFPS system with and without substrates present. It is possible that the introduction of the SAM and theobromine substrates could disrupt the production of TCS1 within the CFPS system. To show that this was not a problem we expressed TCS1 in the CFPS system both with and without the substrates present. There was no notable difference in protein production after the introduction of the substrates, so they did not disrupt protein production.

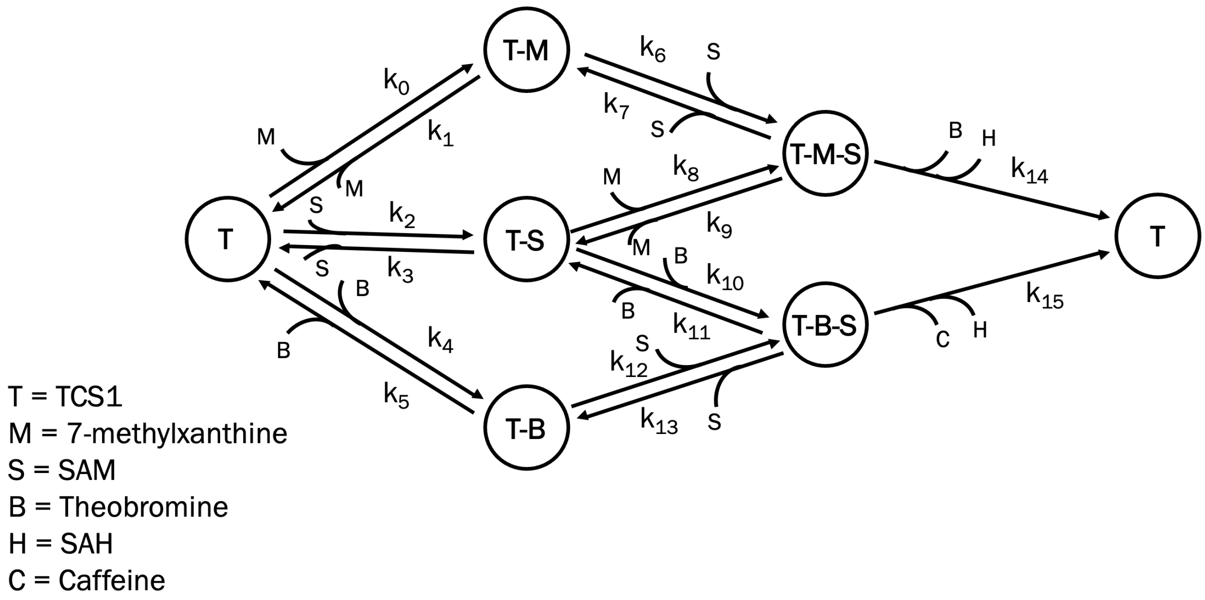


**Figure S2**. A model of the enzyme reactions to go from 7-methylxanthine to theobromine to caffeine. The reactions to methylate 7-methylcanthine 2 additional times can be modeled with enzyme dynamics to better understand what is occurring when TCS1 performs its methylations. In this diagram each node represents a binding state of the enzyme, where T is the TCS1 enzyme alone, and each combination of letters represents the enzyme bound to a different combination of substrates and cofactors. The system feeds back on itself as the theobromine is produced, then the methylation of theobromine competes with the methylation of 7-methylxanthine.

$$dT/dt = -k0*T*M+k1*TM-k2*T*S+k3*TS-k4*T*B+k5*TB+k14*TMS+k15*TBS$$
$$dTM/dt = +k0*T*M-k1*TM-k6*TM*S+k7*TMS$$
$$dTS/dt = +k2*T*S-k3*TS-k8*TS*M+k9*TMS-k10*TS*B+k11*TBS$$
$$dTB/dt = +k4*T*B-k5*TB-k12*TB*S+k13*TBS$$
$$dTMS/dt = +k6*TM*S-k7*TMS+k8*TS*M-k9*TMS-k14*TMS$$
$$dTBS/dt = +k10*TS*B-k11*TBS+k12*TB*S-k13*TBS-k15*TBS$$
$$dM/dt = -k0*T*M-k8*TS*M+k1*TM+k9*TMS+k11*TBS$$
$$dS/dt = -k6*TM*S-k2*T*S-k12*TB*S+k3*TS+k7*TMS+k13*TBS$$
$$dB/dt = -k4*T*B-k10*TS*B+k5*TB+k14*TMS$$
$$dC/dt = +k15*TBS$$
$$dH/dt = +k14*TMS+k15*TBS$$

**Figure S3**. A set of differential equations that can be used to quantitatively simulate the reaction model depicted in Figure S2. Based on the model we constructed in Figure S2 we can simulate the reactions based on a set of constants that represent the rate of each step of the reaction and the concentration of each compound in solution. These mass balance differential equations were then used to observe a simulation of how the reaction proceeds. However, not all the rate limiting steps for this reaction are known, so the simulation has limited accuracy unless they are measured.

The enzyme kinetics model and the differential equations were created with the help of Enzo, a web tool for derivation and evaluation of kinetic models of enzyme catalyzed reactions.[1]

| CONSTANT | VALUE |
| --- | --- |
| **k0** | 80 $\mu M^{-1}s^{-1}$ |
| **k1** | 100 $s^{-1}$ |
| **k2** | 1.14 $\mu M^{-1}s^{-1}$ |
| **k3** | 41 $s^{-1}$ |
| **k4** | 80 $\mu M^{-1}s^{-1}$ |
| **k5** | 100 $s^{-1}$ |
| **k6** | 0.15 $\mu M^{-1}s^{-1}$ |
| **k7** | 1 $s^{-1}$ |
| **k8** | 10 $\mu M^{-1}s^{-1}$ |
| **k9** | 1 $s^{-1}$ |
| **k10** | 10 $\mu M^{-1}s^{-1}$ |
| **k11** | 1 $s^{-1}$ |
| **k12** | 0.15 $\mu M^{-1}s^{-1}$ |
| **k13** | 1 $s^{-1}$ |
| **k14** | 1.5 x $10^{-3}$ |
| **k15** | 3 x $10^{-3}$ |
| **T** | 10 mM |
| **M** | 20 mM |
| **S** | 50 mM |

**Table T1**. The values of each constant used in the simulation. The initial values for T, M, and S are also included, though they can be changed based on the conditions that will be simulated. The rate constants were estimated based on the rate constants found in previous work on other SAM-dependent methyltransferases and are likely only somewhat accurate to the true constants for these reactions. [2,3,4,5]

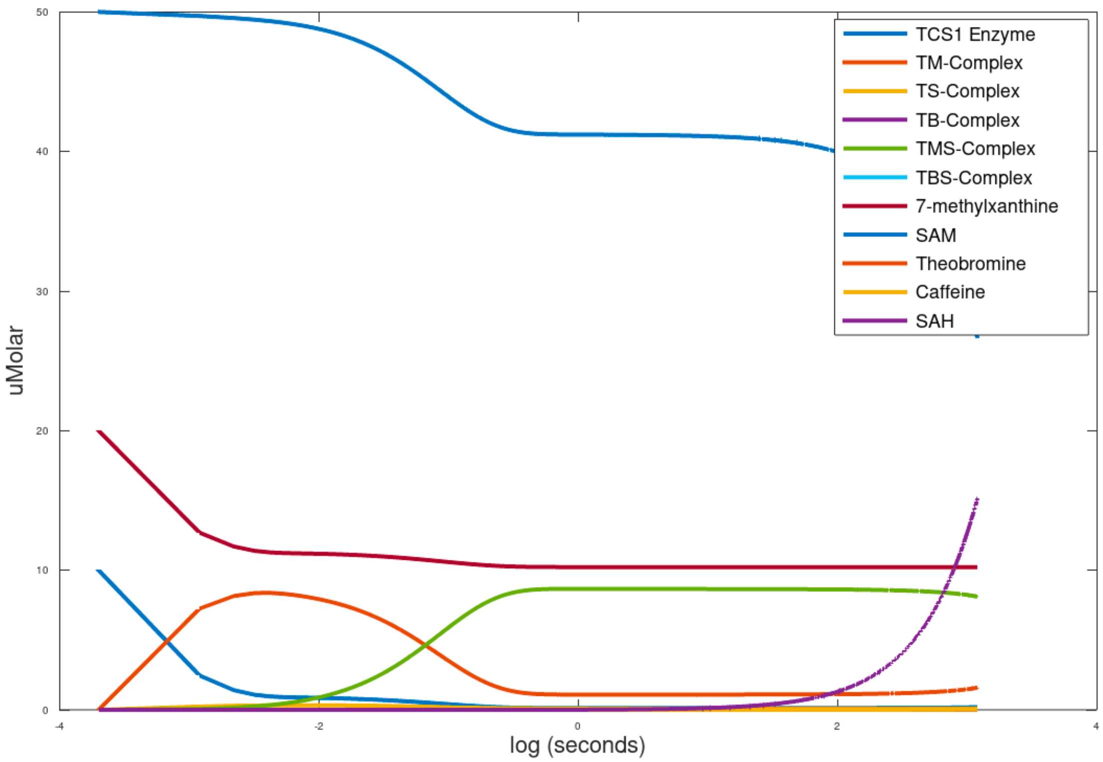


**Figure S4**. A simulation of the methylation reactions going from 7-methylxanthine to theobromine to caffeine. This simulation, based on the model shown in Figure S3, shows the progression of the reactions and the enzyme binding states over a 100 second period on a logarithmic scale. We can observe that the binding of the reactants to the TCS1 enzyme is relatively fast and the rate limiting step is the catalysis of the reaction by TCS1.

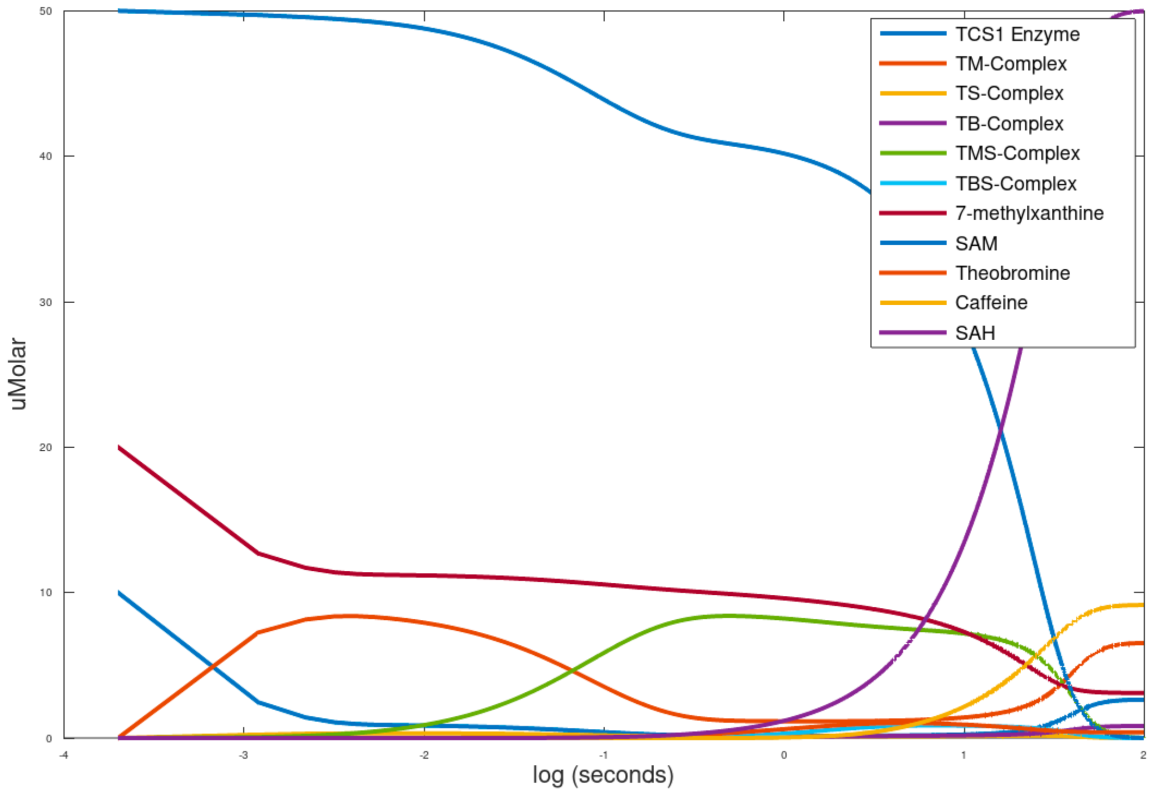


**Figure S5**. A simulation of the methylation reactions going from 7-methylxanthine to theobromine to caffeine with the rate limiting step sped up 100x. This simulation increases the speed of the rate limiting step by increasing the kcat of TCS1 by 100x in order to get a more visually succinct graph without the period between 1 and 100 seconds where the fast parts of the reaction are mostly done and the slow parts had barely started. This simulation is also much faster to run on a computer due to the increased kcat. It shows the reaction going to completion with mostly caffeine produced, but also some theobromine left over and tells us that the time scale for the reaction is on the scale of a few hours. Accuracy is limited by the fact that most of the reaction rate constants were estimated, as they had not directly measured by previous experiments.
